# Altered endothelial mitochondrial Opa1-related fusion in mouse amplifies age-associated vascular and kidney damages

**DOI:** 10.1101/2025.01.29.635602

**Authors:** Carlotta Turnaturi, Loïck L’Hoste, Coralyne Proux, Linda Grimaud, Emilie Vessieres, Antonio Zorzano, Anne Teissier, Pascal Reynier, Raffaella Sorrentino, Guy Lenaers, Laurent Loufrani, Daniel Henrion

## Abstract

**Background:** Cardiovascular diseases are the major cause of death worldwide and their frequency increases with age in association with progressive kidney damages. Endothelial cells (ECs) are early affected in cardiovascular diseases. Although energy production in ECs involves glycolysis, endothelial mitochondria play a role in modulating cellular signalling. A reduction in fusion protein Opa1 level in ECs decreases the vascular response to flow and increased oxidative stress in perfused kidneys. Thus, we hypothesized that reduced Opa1 expression contributes to vascular aging.

**Methods:** We used male and female mice with ECs specific Opa1 knock-out (EC-Opa1), and littermate wild-type (EC-WT) mice aged 6 (young) and 20 months (old). Mesenteric resistance arteries (MRA) and kidneys were collected for vascular reactivity and Western-blot analysis.

**Results:** In old EC-Opa1 mice blood urea was greater than in age-matched EC-WT mice and MRA showed hypercontractilty and reduced endothelium-dependent relaxation. In kidneys, the mitochondria fission protein Fis-1 and the peroxisome proliferator-activated receptor gamma coactivator-1 alpha (Pgc-1α) were increase in old EC-Opa1 mice. The level of eNOS expression was greater in young EC-Opa1 mice and caveolin-1 expression greater in old EC-Opa1 mice. Moreover, in kidneys from EC-Opa1 old mice, NADPH-oxidase subunits gp91, p47 and p67 expression was greater than in age-matched EC-WT mice. No difference was observed between old and young EC-WT mice.

**Conclusion:** Reduced mitochondrial fusion in mouse ECs altered mesenteric vascular reactivity and increased oxidative stress in aging kidneys. Thus, Opa1 might protect the vascular tree in target organs such as the kidney during aging.

## Introduction

Aging is the most important risk factor for cardiovascular disorders ^1,2^. The endothelium has a central role in vascular homeostasis, and endothelial cells (ECs) are the front-line cells against vascular diseases ^3^ explaining why dysregulation of vascular endothelial cells is one of the major cause of cardiovascular disease (CVD) as atherosclerosis, arterial hypertension, coronary artery disease, ischemia reperfusion injury and myocarditis ^4^. The kidney has a dense microvascular network and is early affected as a target organ in CVD. ECs have a small number of mitochondria and their ATP production relies mainly on glycolysis and not on mitochondria ^5^. Nevertheless, mitochondria have a role in the regulation of ECs function ^6^. They participate in cellular homeostasis and have a role in reactive oxygen species (ROS) production and in the regulation of Ca^2+^ concentration in the cytosol ^7^. Mitochondria are dynamic organelles that undergo continuous cycles of fusion and fission. Mitochondrial fusion is mediated by mitofusin-1 (MFN1), mitofusin-2 (MFN2) and optic atrophy type 1 (OPA1) and fission is mediated by dynamin-related protein 1 (DRP1) and mitochondrial fission-1 protein (FIS1). MFN1 and MFN2 are responsible of the outer membrane fusion while OPA1 is necessary for the inner membrane fusion. For fission, DRP1 a cytosolic protein aggregates to FIS1 and anchored to the outermembrane to promote mitochondrial membrane fission. An equilibrated balance between fission and fusion is important for mitochondrial functions. OPA1 is also important for the maintenance of cristae structure of mitochondria and OXPHOS respiration ^8^.

The formation of new mitochondria is regulated by the peroxisome proliferation-activated receptor gamma co-activator 1α (PGC-1α) which is a transcriptional activator of nuclear respiratory factor (Nrf)1. Mitochondrial biogenesis diminishes with age, leading to a mitochondrial dysfunction ^9^. A loss of OPA1 leads to a disorganization of cristae and mitochondrial fragmentation, with mitochondrial dysfunction associated to an excessive ROS production ^10–13^. We have previously shown that *Opa1*^+/-^ mice are more susceptible to hypertension ^14^ and that flow-mediated dilatation (FMD) is selectively reduced in a mouse model with Opa1 deficiency in ECs only (EC-Opa1 mice) ^15^. In this study, FMD was reduced in small resistance arteries to generate a deterioration of the flow-pressure relationship in the perfused kidney, in young EC-Opa1 mice. This was associated with an excess production of ROS in arteries and in the kidney. No difference was observed between male and female mice ^15^.

Thus, we hypothesized that the mitochondrial fusion deficiency in ECs accelerates vascular and kidney damages in aging. To address this question, we used the mouse model previously described with *Opa1* knock-out in ECs ^15^ and investigated vascular reactivity in mesenteric arteries and oxidative stress and inflammation in the kidney in male and female mice aged 6- and 20-months.

## Materials and Methods

### Mice

As previously described ^15^ mice lacking Opa1 in ECs were obtained after crossing *Cadherin5-CreERT2* mice with *Opa1^loxP/loPx^* mice ^16^. They were designed as EC-Opa1 mice (*Cadherin5-CreERT2^+^Opa1^loxP/loPx^*) and were compared to their littermate control EC-WT mice (*Cadherin5-CreERT2^-^Opa1 ^loxP/loxP^*). The deletion was induced by injection of tamoxifen (150 mg/kg per day, diluted in corn oil) during 5 consecutive days in mice aged 3 months. Mice were then used 3 (young mice) or 17 months (old mice) after tamoxifen induction. Thus, mice included in the study were aged 6 months (young mice) or 20 months (old mice). Male and female mice were used in the study. In a previous study using EC-Opa1and EC-WT we have not observed significant differences between male and female mice ^15^.

All procedures were performed in accordance with the principles and guidelines established by the National Institute of Medical Research (INSERM) and were approved by the local Animal Care and Use Committee (APAFIS#2018011217209, APAFIS#30385-2021031010145750). The investigation conforms to the directive 2010/63/EU of the European parliament.

### Vascular reactivity in mesenteric arteries *in vitro*

Segments of first order mesenteric arteries were carefully dissected free of fat and connective tissues. They were then mounted in a 610 M wire-myograph (Danish MyoTechnology, DK) as previously described ^17^. Briefly, two tungsten wires were inserted into a 2 mm long arterial segment; one was fixed to a force transducer and one to a micrometer. They were continuously bathed in a physiological salt solution (PSS) of the following composition (mM): 130, NaCl; 15, NaHCO_3_; 3.7, KCl; 1.2 KH_2_PO_4_; 1.2, MgSO_4_; 11, glucose; 1.6, CaCl_2_; and 5, HEPES, pH 7.4, pO_2_ 160 mmHg, pCO_2_ 37 mmHg. Wall tension was applied as described previously ^18^. Arterial contractility was tested using phenylephrine (Phe, 10^-9^ to 3.10^-5^ mol/L). Endothelial function was then tested using acetylcholine (Ach, 10^-9^ to 3.10^-5^ mol/L) after precontraction with Phe (10^-6^ mol/L). Endothelium-independent relaxation was tested with sodium nitroprusside (SNP, 10^-9^ to 10^-5^ mol/L) after precontraction with Phe (10^-6^ mol/L).

### Analysis of protein expression levels by Western blot

Kidney proteins were extracted in extraction buffer : SDS 0,1%, Tris 10 Mm pH 7,4, proteases inhibitors 1X (CAT#78444, Thermo Fisher Scientific, Waltham, MA, USA), EDTA 0,5 mM. Homogenates were centrifuged at 13000 rpm at 4°C for 20 min, and the resulting supernatant was collected. Protein concentration was determined using Micro BCA protein assay kit (cat#23227, Thermo Fisher Scientific, Waltham, MA, USA) according to the manufacturer’s instructions. Equal amounts of proteins (30 µg) were solubilized in 25 µl of Laemmli sample buffer containing 2,5% β-mercaptoethanol, boiled 5 min at 90°C, separated by 4-15% polyacrylamide gel electrophoresis (BioRad, Marnes la Coquette, France) and transferred to a nitrocellulose membrane (BioRad). Membranes were incubated overnight at 4 °C with the primary antibody followed by the appropriate peroxidase-labeled secondary antibody (see the Major Resources Table in the supplement files) for 1 h. Reactions were visualized by ECL detection according to the manufacturer’s instructions (Bio-Rad, Marnes-la-Coquette, France) and membranes were stripped at room temperature 20 minutes twice in the presence of a low pH glycine solution before re-blotting.

### Blood urea nitrogen level in mice

Blood urea nitrogen was measured using the Atellica™ CH Urea Nitrogen (UN-c) assay (Siemens, Erlangen, Germany).

### Statistical analyses

For concentration-response curves, a two-way ANOVA for repeated measurements followed by a Bonferroni’s post-test was performed. For the other comparisons, 2-way ANOVA followed by a Bonferroni’s post-test was used, as indicated in the figure legends. Probability values lower than 0.05 were considered significant. Data from males and females mice were pooled.

## Results

### Blood urea nitrogen measurements

Blood urea nitrogen was equivalent in young EC-Opa1 and EC-WT (Figure 1). It was not significantly different in old EC-Opa1 mice compared to young EC-WT mice (Figure 1). Nevertheless, blood urea nitrogen was significantly elevated in old EC-Opa1 mice compared to old EC-WT mice (Figure 1).

**Figure 1.**
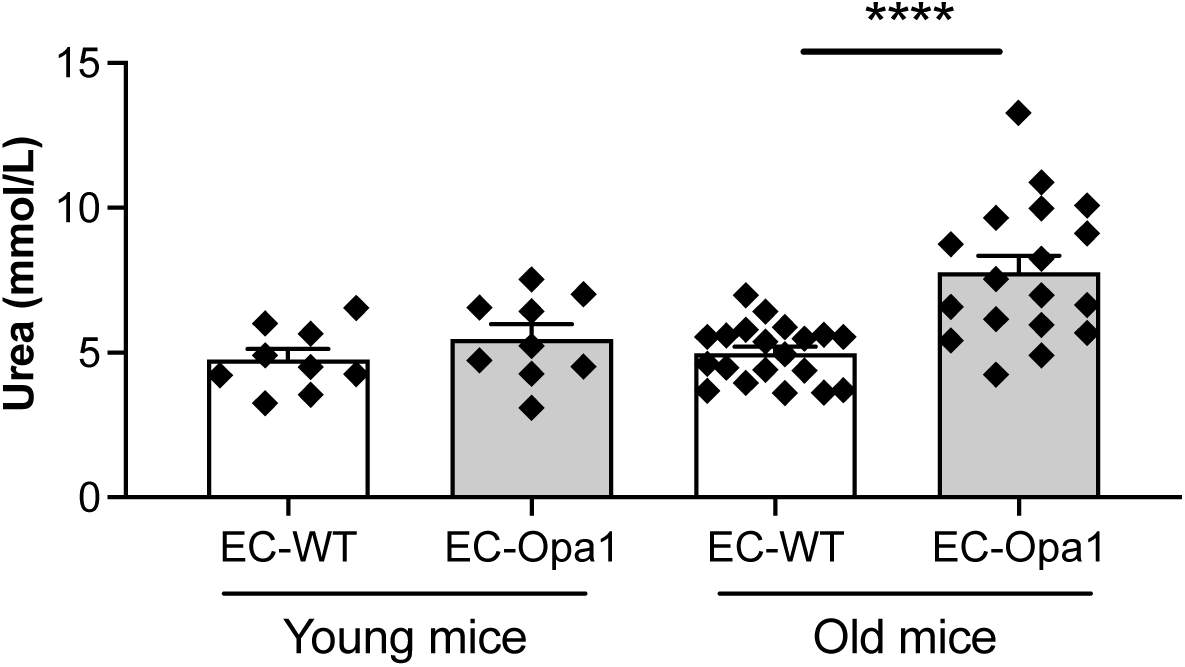
Blood urea nitrogen measurement. Blood urea nitrogen was measured in young and old EC-Opa1 and EC-WT mice. Data is expressed as means ± SEM (n=9 EC-WT mice, 9 young EC-Opa1 young mice, 20 old EC-WT mice and 18 EC-Opa1 old mice). ****p<0.0001, two-way ANOVA and Bonferroni’s multiple comparisons test.

### Vascular contractility in mesenteric arteries

Phenylephrine-mediated contraction was measured in isolated mesenteric arteries. In both young and old mice, phenylephrine-mediated contraction was significantly higher in EC-Opa1 than in age mached EC-WT mice (figure 2A,B).

**Figure 2:**
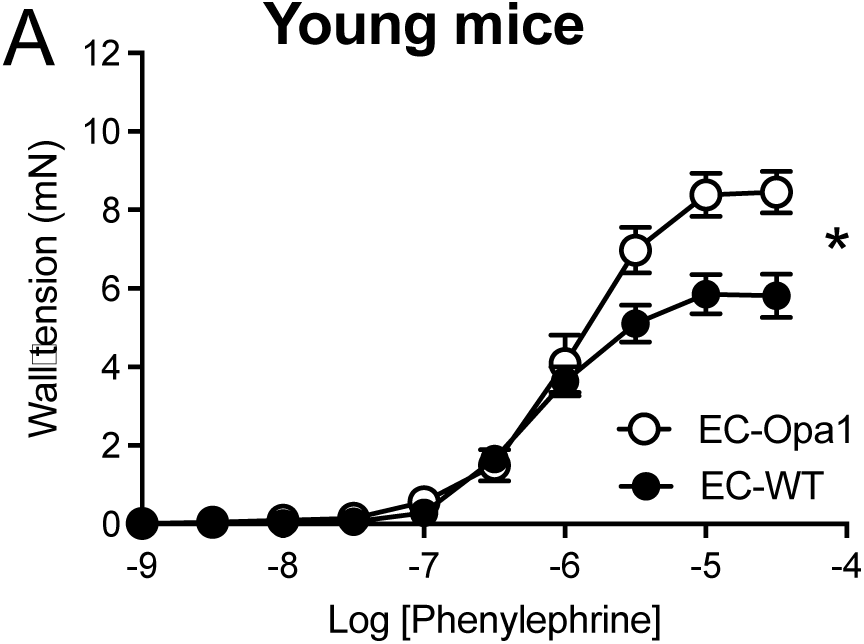
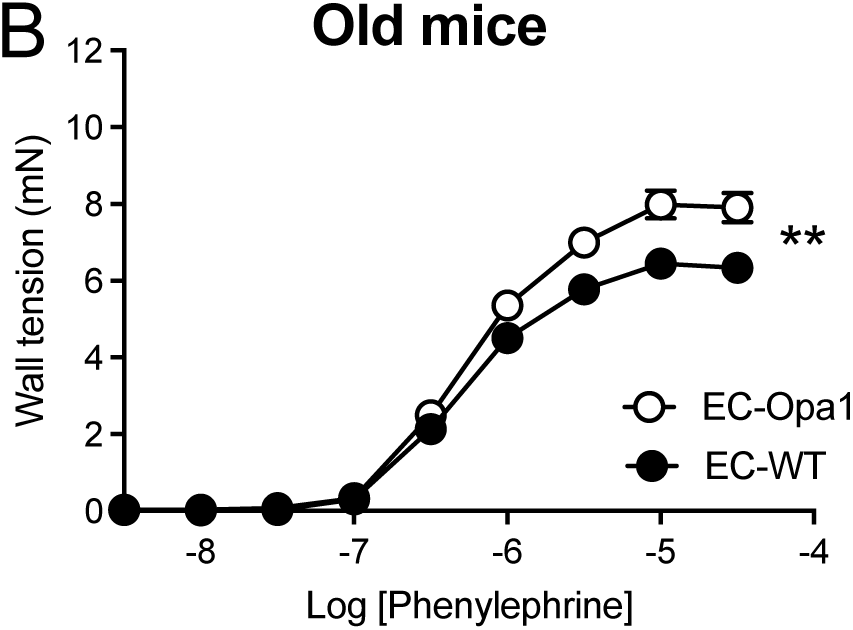
Phenylephrine-mediated contraction. Phenylephrine (1 nmol/L to 30 µmol/L)-mediated cumulative concentration-response curve was determined in mesenteric arteries isolated from young and old EC-Opa1 and EC-WT mice. Data is expressed as mean ± SEM (n=12 young EC-Opa1 mice, 13 young EC-WT mice, 19 old EC-Opa1 mice and 21 old EC-WT mice). *p<0.05, **p< 0.01, two-way ANOVA for repeated measurements and Bonferroni’s multiple comparisons test.

### Endothelium-dependent relaxation in mesenteric arteries

Acetylcholine (Ach) induced endothelium-dependent relaxation in mesenteric arteries (Figure 3). In young mice ACh-mediated relaxation was equivalent in EC-Opa1 and EC-WT mice (figure 3A) while in old mice a significant reduction in relaxation was observed in EC-Opa1 mice compared to EC-WT mice (figure 3B).

**Figure 3:**
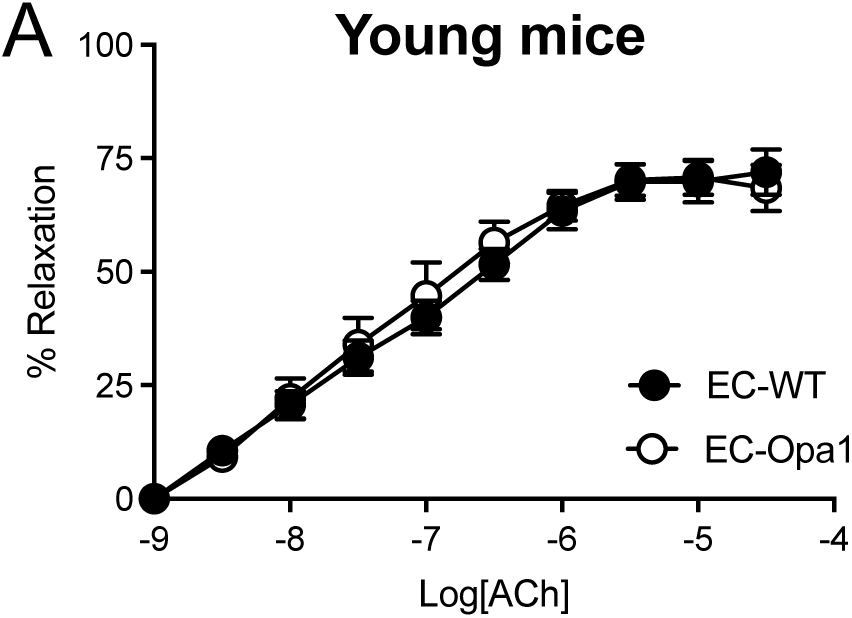
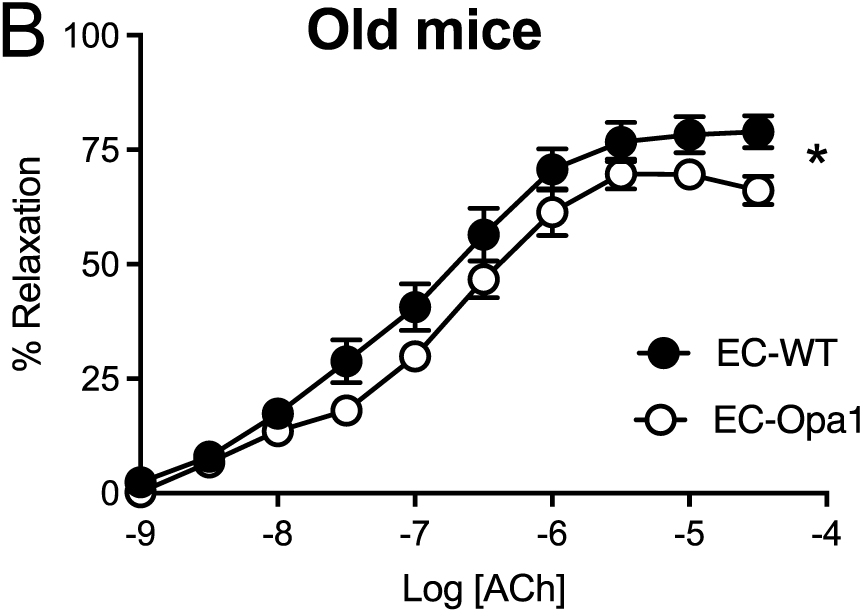
Acetylcholine-mediated endothelium-dependent relaxation. Acetylcholine (Ach, 1 nmol/L to 30 µmol/L)-mediated endothelium-dependent relaxation was determined in mesenteric arteries isolated from young and old EC-Opa1 and EC-WT mice. Data is expressed as means ± SEM (n=12 young EC-Opa1 mice, 13 young EC-WT mice, 19 old EC-Opa1 mice and 21 old EC-WT mice). *p<0.05, two-way ANOVA for repeated measurements and Bonferroni’s multiple comparisons test.

### Endothelium-independent relaxation in mesenteric arteries

Sodium nitroprusside (SNP) induced endothelium-independent relaxation in isolated mesenteric arteries (Figure 4). We observed no significant difference between groups.

**Figure 4:**
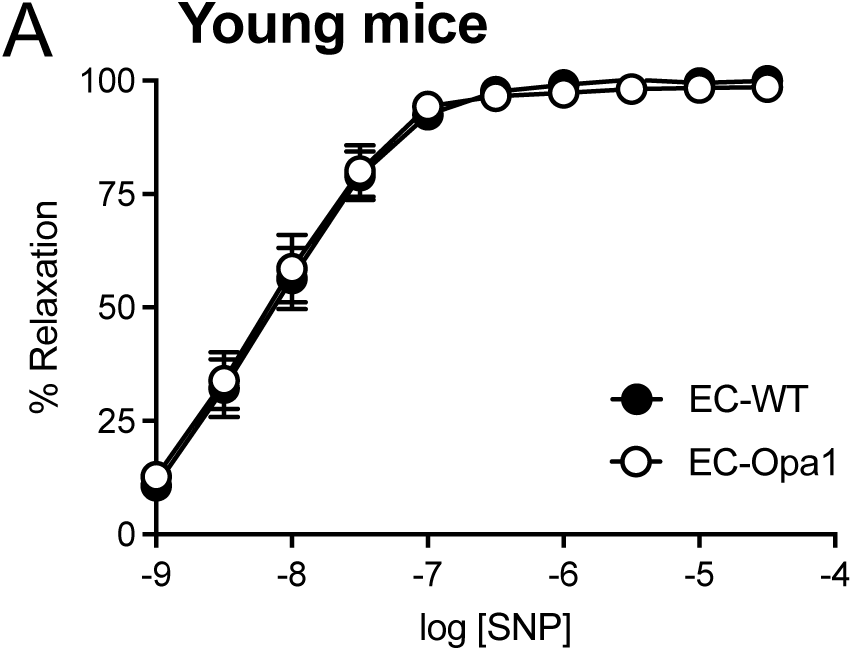
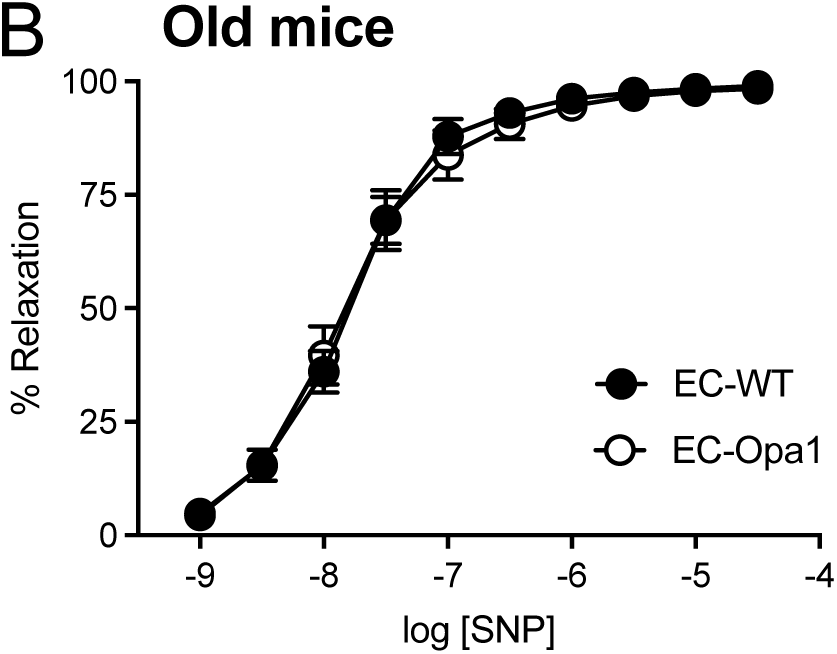
Sodium nitroprusside-mediated endothelium-independent relaxation. Sodium nitroprusside (SNP, 1 nmol/L to 30 µmol/L)-mediated endothelium-dependent relaxation was determined in mesenteric arteries isolated from young and old EC-Opa1 and EC-WT mice. Data is expressed as means ± SEM (n=12 young EC-Opa1 mice, 13 young EC-WT mice, 19 old EC-Opa1 mice and 21 old EC-WT mice). NS, Two-way ANOVA for repeated measurements and Bonferroni’s multiple comparisons test.

### Analysis of protein expression in kidneys

The expression level of the mitochondrial fission protein Fis1 was significantly increased in the kidney in old EC-Opa1 mice compared to old EC-WT mice (figure 5A) while the expression level of the mitochondrial fusion protein Mfn2 was not affected (figure 5B). The expression level of Fis-1 and Mfn2 was not affected by the absence of Opa1in the endothelium in young mice (Figure 5A and B).

**Figure 5:**
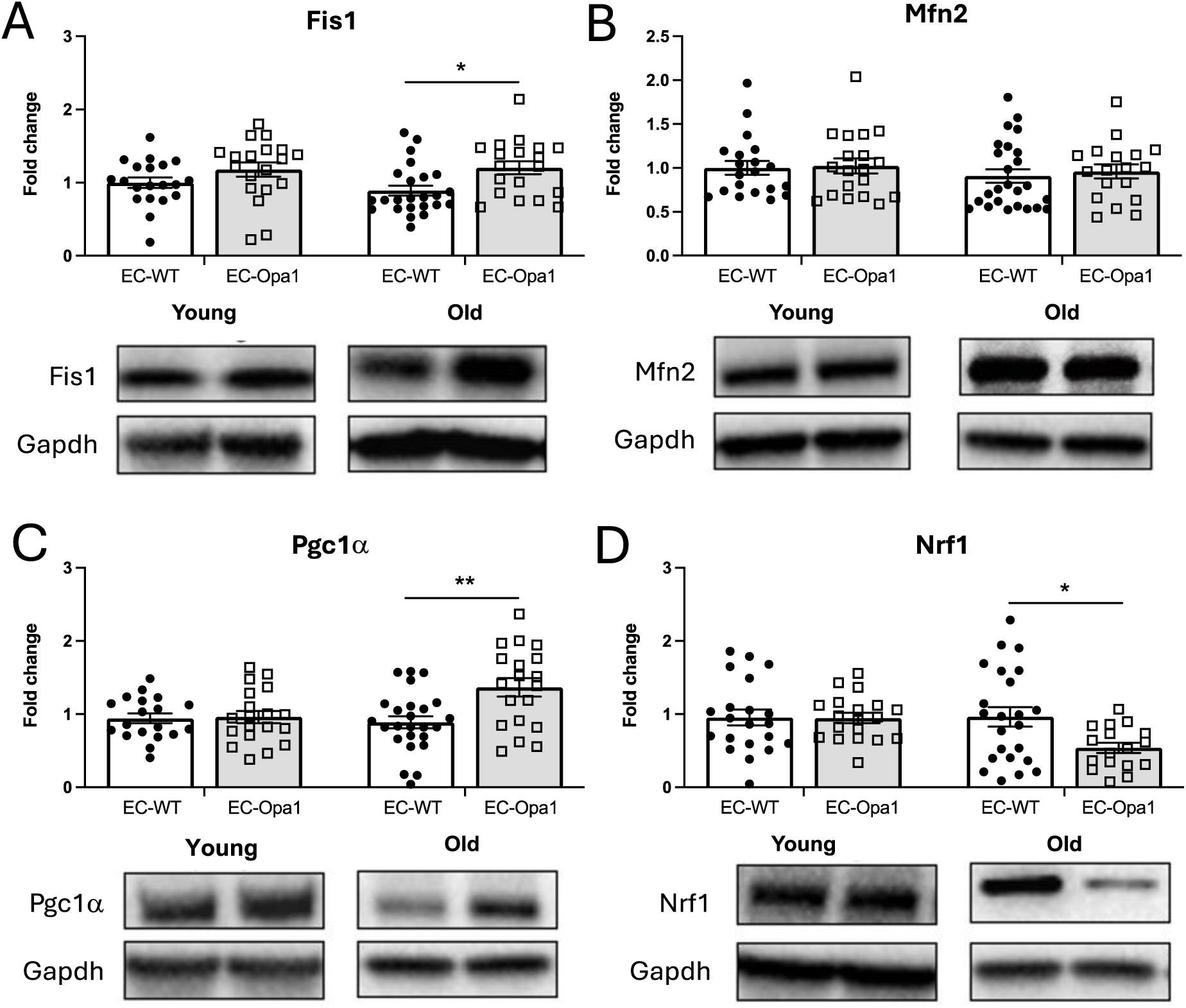
Protein expression level of Fis1, Pgc1α, Mfn2 and Nrf-2 in the kidney. Protein expression level of Fis1 (A), Mfn2 (B), Pgc1α (C) and Nrf-1 (D) was determined in kidneys isolated from young and old EC-WT and EC-Opa1. Mean±SEM is shown (n=21 to 27 mice per group). Uncropped blots are shown as supplemental files. *p<0.05 and **p<0.01, two-way ANOVA and Bonferroni’s multiple comparisons test.

The expression level of Pgc-1α, which promotes mitochondrial biogenesis, was significantly increased in old EC-Opa1 mice compared to old EC-WT mice (figure 5C), while no difference was observed in young mice. Surprisingly, Nrf-1 expression level, which is transcriptionally activated by Pgc-1α, was significantly decreased in EC-Opa1 old mice compared to EC-WT old mice (figure 5D).

The expression level of eNOs in kidneys was significantly increased in young EC-Opa1 mice compared to young EC-WT mice (figure 6A) whereas no significant change in eNOs expression level was observed in old mice EC-Opa1 old mice compared to old EC-WT mice.

**Figure 6:**
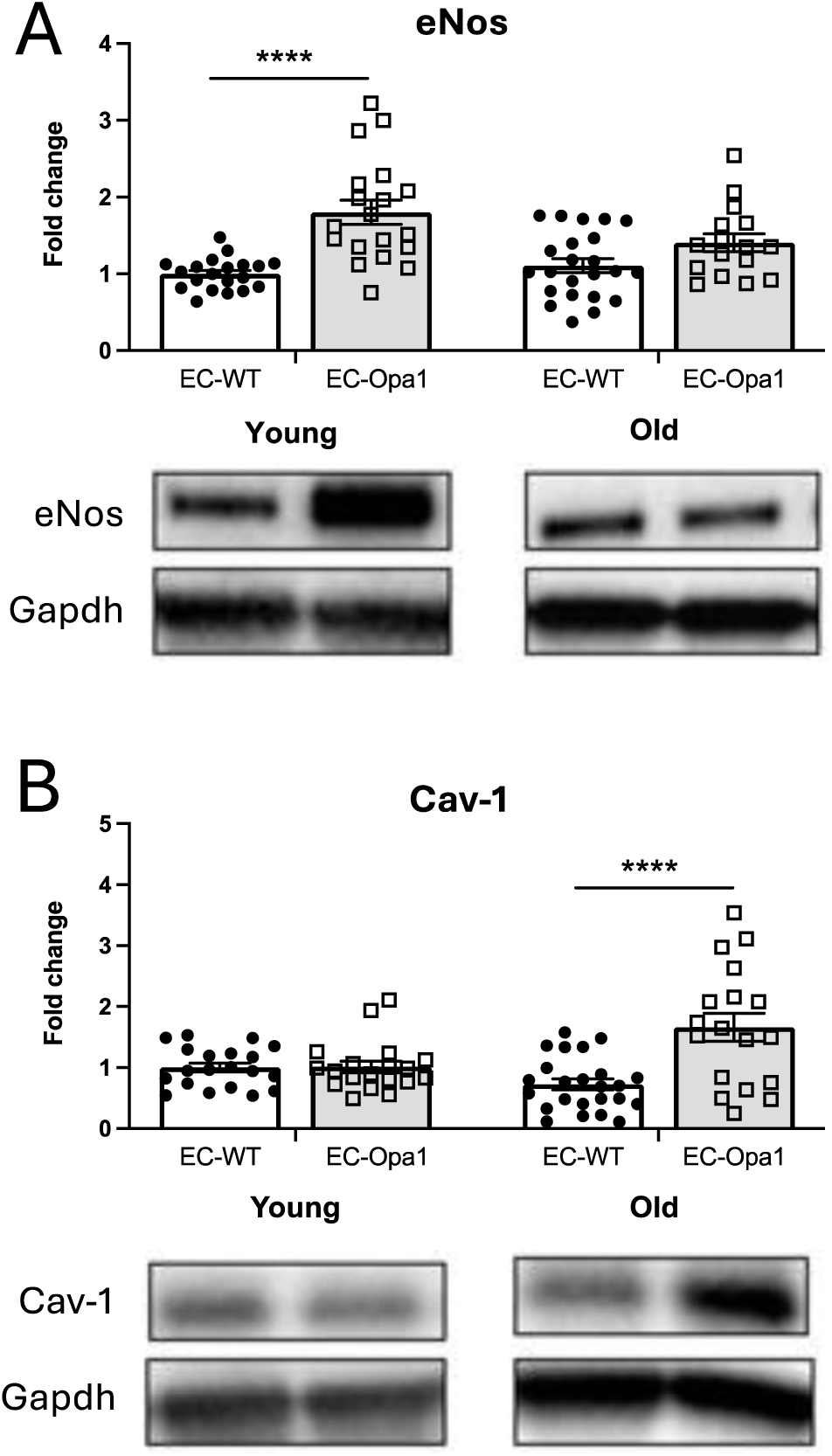
Protein expression level of eNos and Cav-1 in the kidney. Protein expression level of eNos (A) and Cav-1 (B) was determined in kidneys isolated from young and old EC-WT and EC-Opa1 mice. Mean±SEM is shown (n= 21 to 27 mice per group). Uncropped blots are shown as supplemental files. ****p<0.0001, two-way ANOVA and and Bonferroni’s multiple comparisons test.

By contrast, Cav-1 expression level was significantly increased in old EC-Opa1 mice compared to old EC-WT mice without significant change in young mice (figure 6B).

The expression level of Gp91^phox^ in kidneys was significantly increased in old EC-Opa1 mice compared to old EC-WT mice (figure 7A) whereas no significant change in Gp91^phox^ expression level was observed in young mice EC-Opa1 mice compared to young EC-WT mice.

**Figure 7:**
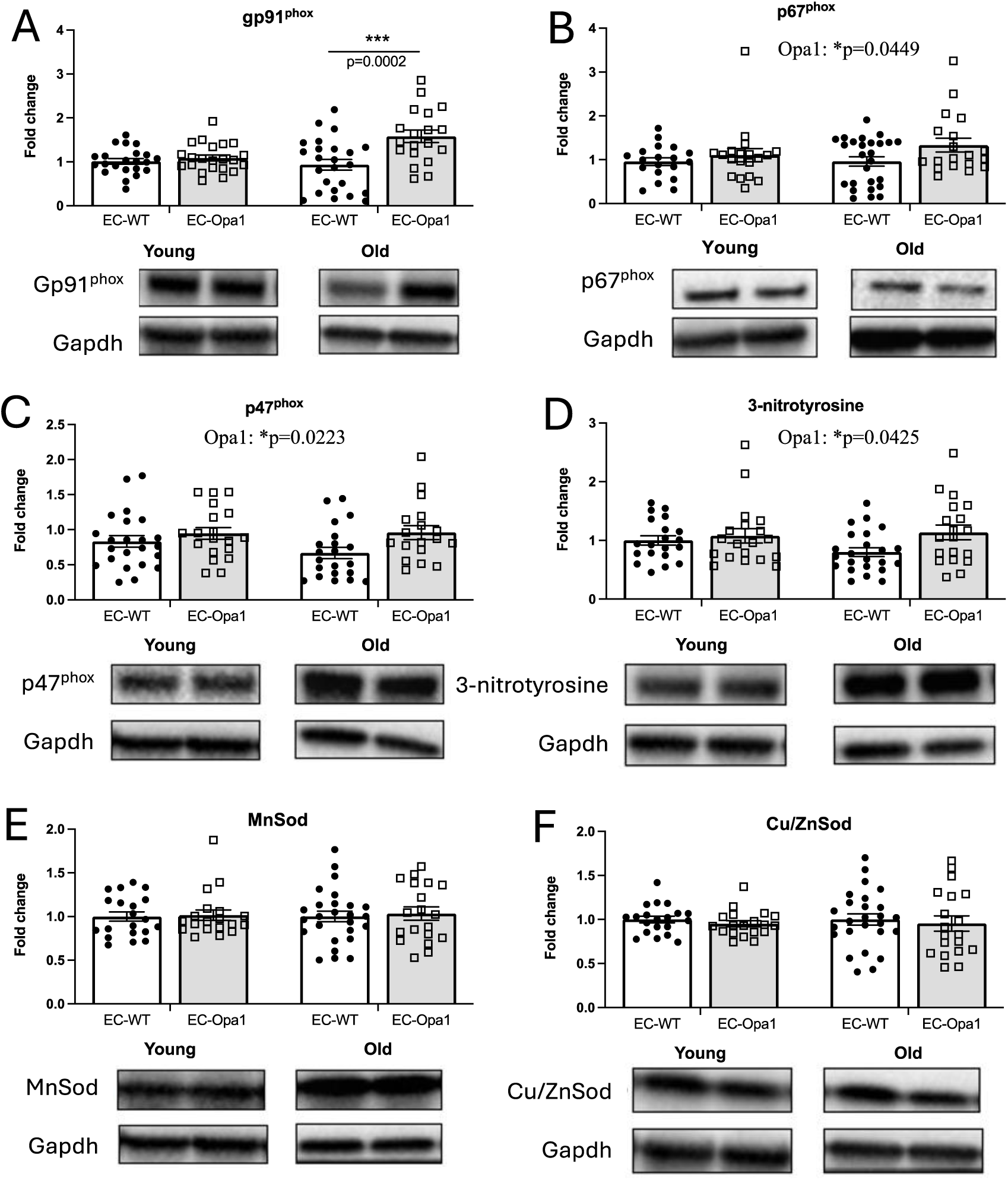
Protein expression levels of Gp91^phox^, p67 ^phox^, p47 ^phox^, 3-nitrotyrosine, MnSod and Cu/ZnSod in the kidney. Protein expression levels of gp91^phox^ (A), p67^phox^ (B), p47 ^phox^ (C), 3-nitrotyrosine (D), MnSod (E) and Cu/ZnSod (F) was determined in kidneys isolated from young and old EC-WT and EC-Opa1 mice. Mean±SEM is shown (n=21 to 27 mice per group). Uncropped blots are shown as supplemental files. Detail of the two-way ANOVA and Bonferroni’s multiple comparisons test: Gp91 **^phox^**: absence of Opa1: ***p=0.0008, age: *p=0.0488, interaction: **p=0.0070, old EC-Opa1 versus old EC-WT mice: ***p=0.0002 (Bonferroni’s multiple comparisons test). p67 **^phox^**: absence of Opa1: *p=0.0449, age: NS, interaction: NS, Bonferroni’s multiple comparisons test: NS. p47 **^phox^**: absence of Opa1: *p=0.0223, age: NS, interaction: NS, Bonferroni’s multiple comparisons test: NS. 3-nitrotyrosine: absence of Opa1: *p=0.0425, age: NS, interaction: NS, Bonferroni’s multiple comparisons test: NS.

The ANOVA analysis identified significant effect of the absence of Opa1 in p47^phox^, p67^phox^ and 3-NT expression level, without significant effect of age (Figure 7B, C and D)

No significant difference between groups was found for MnSod and Cu/ZnSod (figure 7E,F).

## Discussion

The present study shows that reduced Opa1 expression in ECs induces hypercontractility and reduces endothelium dependent relaxation in resistance arteries in old mice. These observations were associated with hyperuremia and oxidative stress in kidneys.

We used 20-months old mice, not older, to avoid the occurrence of important changes in vascular reactivity and kidney function assuming that changes due to the reduced expression in Opa1 in ECs would be more difficult to observe in very old mice with too many organ dysfunctions. Indeed, 20-months old EC-WT mice had no significant change in vascular reactivity and no obvious alteration in kidney function as shown by the absence of difference between young and old EC-WT mice. In these conditions, changes observed in old EC-Opa1 mice are mainly attributed to the reduced expression in Opa1 in ECs.

In a previous study we have shown that in this model (EC-Opa1 mice), Opa1 protein expression is reduced by 61% in ECs ^15^. In this study flow-mediated dilation was reduced in both mesenteric resistance arteries and in isolated perfused kidneys. This was associated with increased oxidative stress. No difference was observed between male and female mice ^15^. Thus, in the present study we pooled male and female mice.

Kidney disease is a major public health problem worldwide and its importance is increasing ^19^. The kidney has one of the richest ECs population linked to a dense microcirculation. The kidney is also a target organ very early affected in CVD ^20^. Thus our results showing reduced endothelium-dependent relaxation and oxidative stress in the kidney on 20-months old EC-Opa1 mice support the assumption that changes in endothelial function affects early kidney function.

Although ECs mitochondria have a restricted role in energy production, as ECs use mainly glycolysis to produce ATP, they take part to ECs function. Endothelial Opa1 has a role in ECs responsiveness to flow (shear stress) as shown by our previous work ^15^ and is necessary for angiogenesis ^21^. Furthermore, reduced Opa1 expression enhanced angiotensin II-induced hypertension in mice ^14^. In old EC-Opa1 mice we found a stronger alteration in vascular reactivity with hypercontractility and reduced endothelium-dependent relaxation. In young EC-Opa1, mice this was not observed, in agreement with our previous work ^15^ showing only a reduction in flow-mediated dilation. Interestingly, it has been demonstrated a significant decrease in Opa1 expression in coronary endothelial cells from a type 1 diabetes mouse model, possibility contributing excessive mitochondrial fission and mitochondrial ROS production ^22^.

Here, we found that in the kidney of old EC-Opa1 mice there is increased Cav-1 expression level. This agrees with a previous work showing that ROS formation increases Cav-1 expression ^23^. Thus, in the present study, Cav-1 expression increased in old EC-Opa1 mice kidneys possibly as a consequence of ROS formation. Furthermore, Cav-1 bound eNOS protein leading to its inactivation and thus, the increased Cav-1 expression could take part to the reduction in endothelium-mediated relaxation observed in old EC-Opa1 mice. In addition, an increase in Cav-1 expression is correlated to acute kidney disease ^24^. This is in line with our results showing an increase in blood urea in aged EC-Opa1 mice. Of note, targeting Cav-1 is also a promising therapeutic route in chronic kidney disease ^24^.

In EC-Opa1 aged kidney, as a consequence of reduction of Opa1 expression, we observed a significant increase of the fission protein Fis1 without any change in Mfn2, the other protein involved in mitochondrial network, suggesting an absence of compensatory mechanism with a possible increase in mitochondrial fission. Fis1 is involved in mitochondrial fission, which produces smaller mitochondria destined for mitophagy ^25^. Our findings showing excessive oxidative stress and vascular disorders in mice lacking ECs Opa1 agrees with previous studies showing that a desequilibrium between fusion and fission occurs in cardiovascular diseases ^6,26–28^.

In addition, we observed a significant increase in Pgc-1α expression in the kidney in old EC-Opa1 mice. Pgc-1α is a major regulator of mitochondrial biogenesis and the reduced OPA1 expression in EC-Opa1 mice could induce an increase in Pgc-1α as a compensatory mechanism. Indeed, increasing Pgc-1α expression in kidney tubular cells improves energy production and protects the kidney whereas increasing Pgc-1α level in ECs deteriorates endothelial function ^29^. The results of the present study do not allow discerning ECs from other cells in the kidney as the whole kidney was used for the western-blot analysis. Thus, we can speculate that the compensatory mechanism is not protective enough to prevent oxidative stress in EC-Opa1 mice. Nevertheless, mitochondrial biogenesis, the formation of new functional mitochondria, is an important defense system for cells to overcome mitochondrial damages ^30^ and Pgc-1α could also be activated by external stimuli such as ROS ^31^. Surprinsly this increase in Pgc-1α was not followed by an increase of Nrf-1 but its expression level decreased, suggesting a possible deregulation in the activation chain of genes that are important for normal mitochondrial functionning. This issue remains to be further explored.

Furthermore, the absence of endothelial Opa1 in old EC-Opa1 mice kidneys was associated with an increased expression of membrane components of NADPH oxidases (p47, p67 and Gp91) suggesting higher ROS formation. This was not followed by an increase in Sod expression, as we did not disclose change in MnSod and Cu/ZnSod expression. These findings suggest that the absence of Opa1 protein in aged kidney led to oxidative stress without a compensatory protective action by the SOD. This increased expression in NADPH oxidase subunits was associated to increased 3-nitrotyrosine levels in the kidneys of EC-Opa1 mice, thus confirming the occurrence of oxidative stress. Increased 3-nitrotyrosine has been previously observed in aged and diabetic mice ^32^. Indeed, protein tyrosine nitration is a good marker of oxidative stress leading to alteration of the activity of the nitrated proteins ^33^. Our observations strongly agrees with our previous work showing increased oxidative stress and inflammation in the kidney of old mice ^17^ and with a previous study showing that silencing Fis1 or Drp1 reduced high glucose-induced alteration in mitochondrial ROS production, indicating that increasing mitochondrial fission could be negative for endothelial function due to an increase in ROS production ^34^.

In conclusion, we demonstrate that endothelial Opa1 has a protective effect during aging by maintaining vascular and kidney function possibly through the reduction of age-associated oxidative stress.

These findings suggest that improving mitochondrial fusion or more generally mitochondrial dynamics could propose a new targets for therapeutic approach against endothelial disorders related to aging and might protect the vascular tree in target organ such as the kidney.

## Data Availability Statement

The data presented in this study are available on request from the corresponding author.

## Acknowledgments

Dr Carlotta Turnaturi was supported by the found: Course of PhD in Pharmaceutical Science, XXXV Cycle, Department of Pharmacy, School of Medicine and Surgery, University of Naples Federico II, Via D. Montesano 49, 80131, Naples, Italy.

The authors thank the animal facility (SCAHU, “Service Commun d’Animalerie Hospitalier et Universitaire”, Angers, France).

## Supplementary Materials

Uncropped blots for the Western-blots shown in figures 5, 6 and 7.

## Major Resources Table

In order to allow validation and replication of experiments, all essential research materials listed in the Methods should be included in the Major Resources Table below. Authors are encouraged to use public repositories for protocols, data, code, and other materials and provide persistent identifiers and/or links to repositories when available. Authors may add or delete rows as needed.

### Genetically Modified Animals

|  | Species | Vendor or Source | Background Strain | Other Information | Persistent ID / URL |
| --- | --- | --- | --- | --- | --- |
| Parent - VeCadh-CreERT2 | Mice | UMR<br>CNRS6015<br>INSERM 1083<br>Angers Dr<br>Henrion | C57Bl6 J |  | MGI:3848982 |
| Parent OPA1flox/flox | Mice | Institute for<br>Research in<br>Biomedicine<br>Barcelona<br>Pr Zorzano | C57Bl6N |  | MGI:6117129 |

### Antibodies

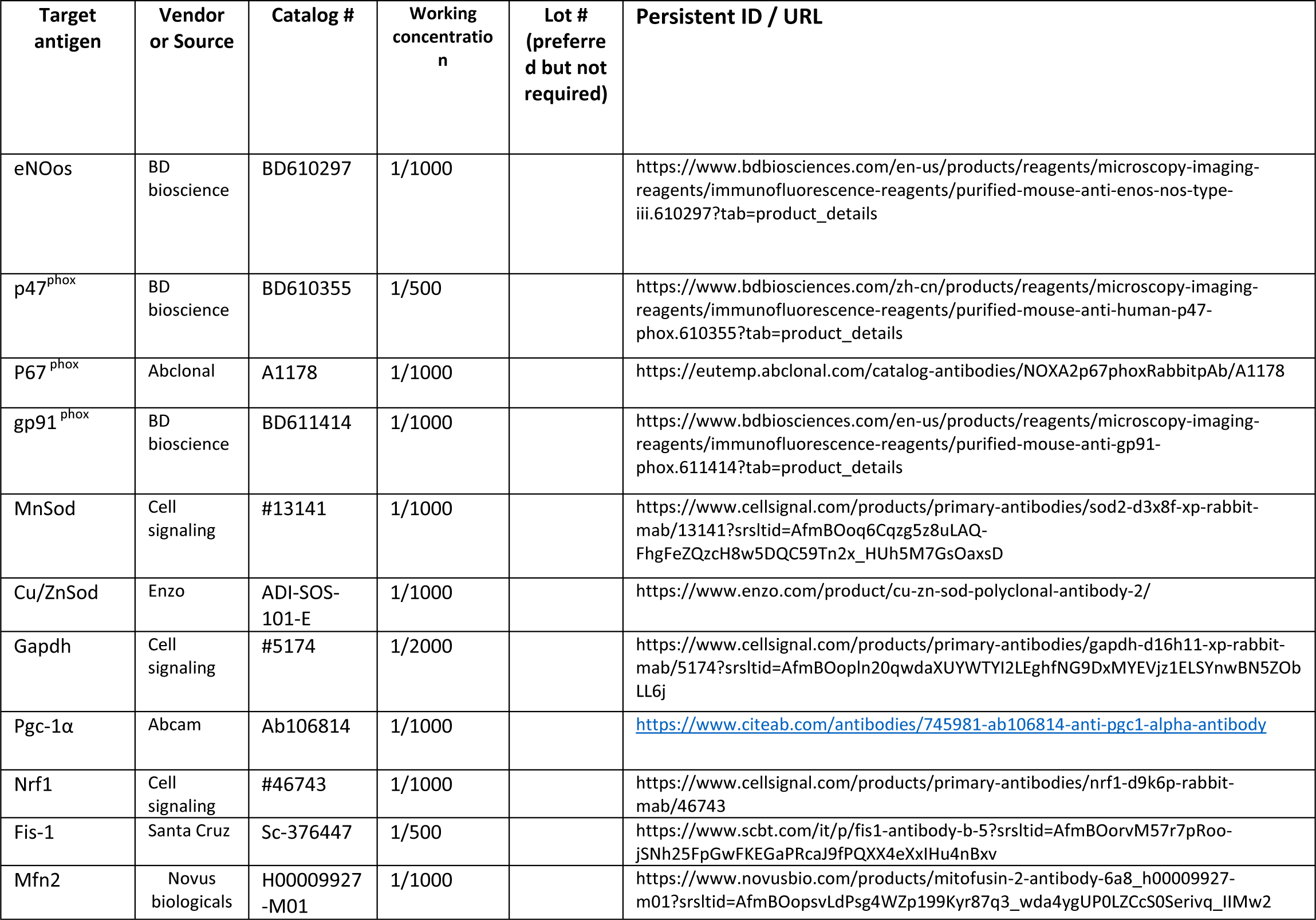
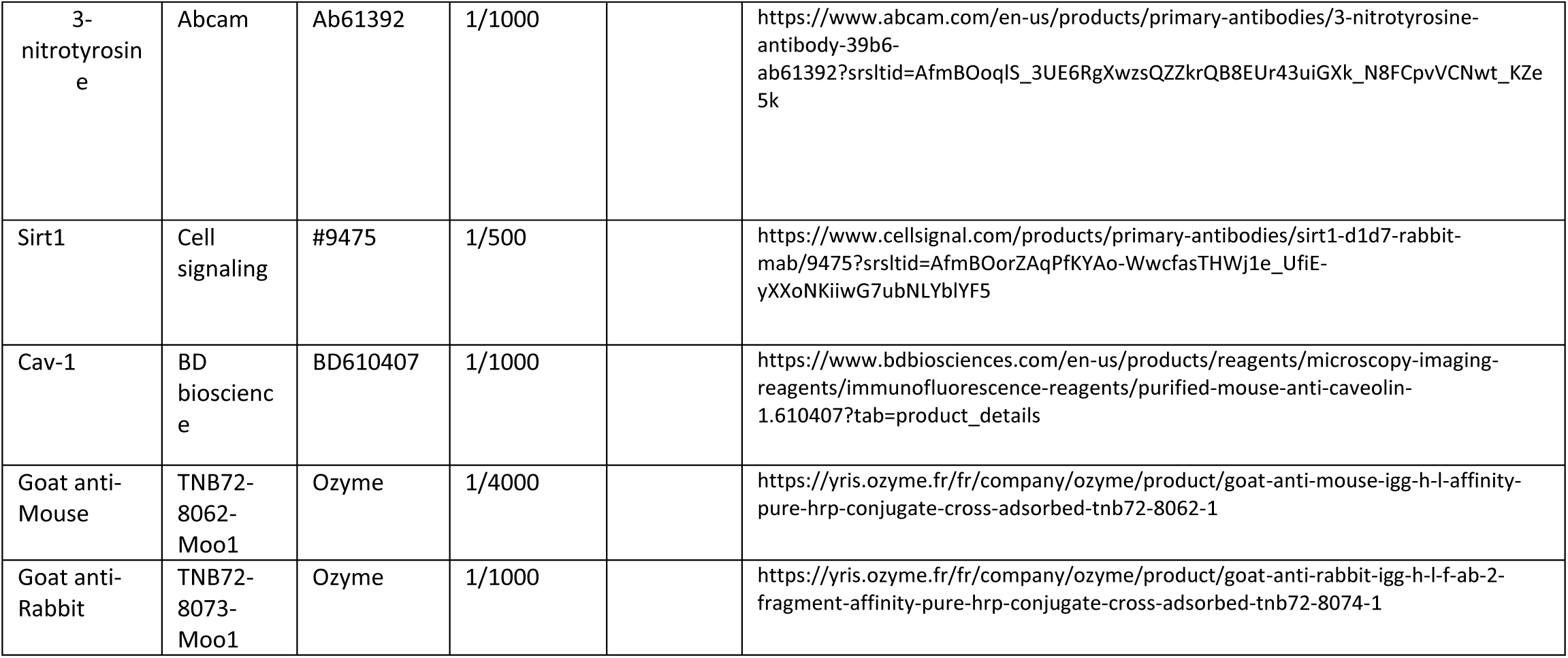

### ARRIVE GUIDELINES

The ARRIVE guidelines (https://arriveguidelines.org/) are a checklist of recommendations to improve the reporting of research involving animals. Key elements of the study design should be included below to better enable readers to scrutinize the research adequately, evaluate its methodological rigor, and reproduce the methods or findings.

#### Study Design

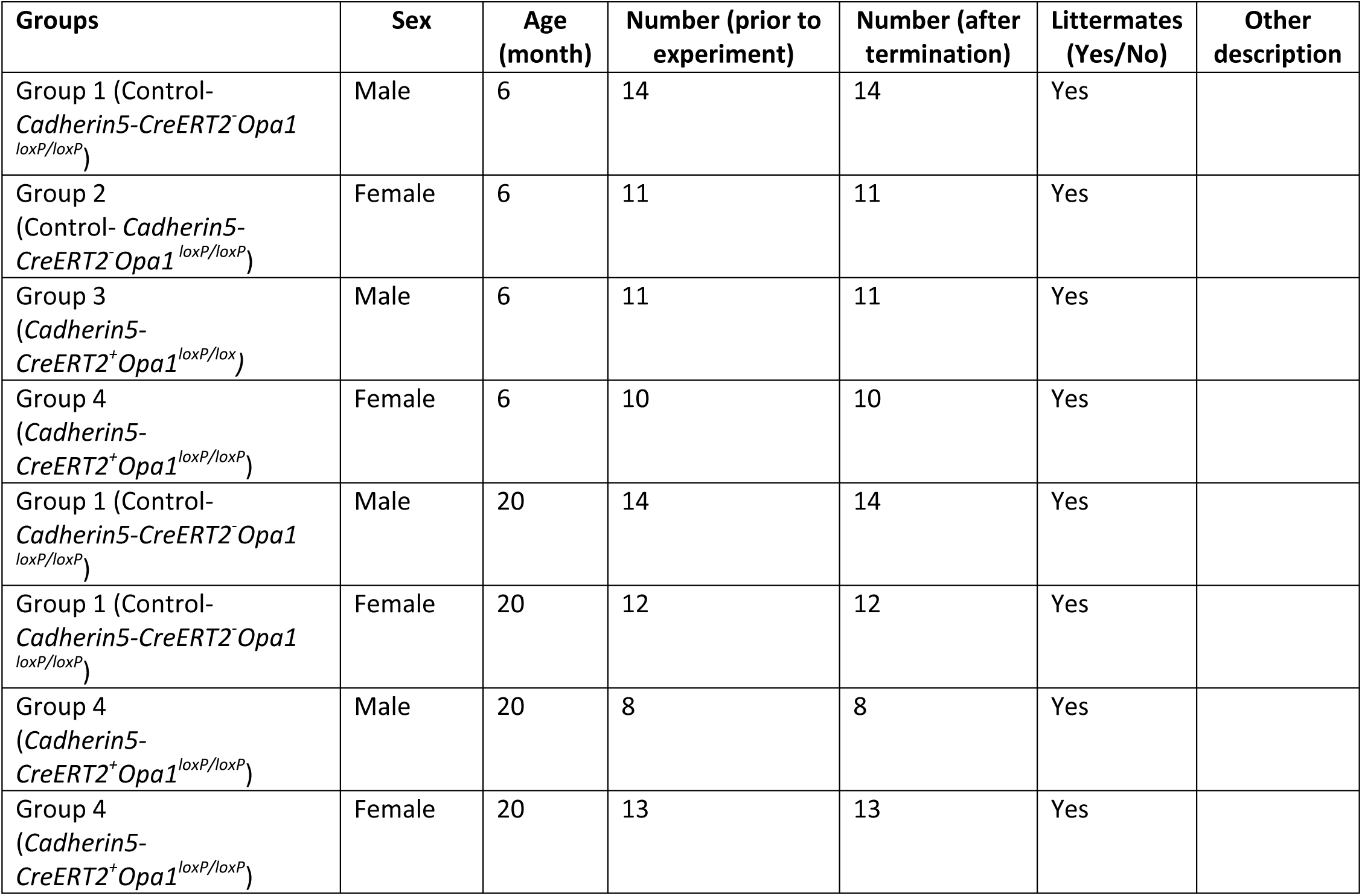

#### Sample Size

Sample size was determined based on our previous studies on the same mice.

#### Inclusion Criteria

Viable and healthy mice born at the University of Angers animal facility (SCAHU).

#### Exclusion Criteria

Exclusion criteria were pre-established to exclude animals that suffer from common mouse abnormality or excessive loss of weight (<20%)

In the present study no animal was excluded.

#### Randomization

Mice were randomly assigned to control (normal diet) or diabetogenic diet.

#### Blinding

Researchers were blinded to genotype or condition during data collection and analysis

